# Social spread of positive facial signals: affiliative zygomatic synchrony in co-present individuals

**DOI:** 10.1101/484980

**Authors:** Golland Yulia, Dana Mevorach, Nava Levit-Binnun

**Author notes:** Corresponding Author details: Yulia Golland, PhD Sagol Center for Brain and Mind Baruch Ivcher School of Psychology Interdisciplinary Center (IDC) Herzliya P.O Box 167 Herzliya, 41650, Israel 972-52-4567168 (voice) 972-9-9602845 (fax).

## Abstract

In social contexts individuals frequently act as social chameleons, synchronizing their responses with those of others. Such synchrony is believed to play an important role, promoting mutual emotional and social states. However, synchrony in facial signals, which serve as the main communicative channel between people, has not been systematically studied. To address this gap, we investigated the social spread of smiling dynamics in a naturalistic social setting and assessed its affiliative function. We also studied whether smiling synchrony between people is linked with convergence in their autonomic and emotional responses. To that aim we measured moment-by-moment changes in zygomatic electromyography and cardiovascular activity in dyads of previously unacquainted participants, who co-viewed emotional movies together. We found a robust, dyad-specific, zygomatic synchrony in co-viewing participants, after controlling for movie-driven effects. During the positive movie, zygomatic synchrony co-varied with cardiovascular synchrony and with convergence in positive feelings. No such links were found for the negative movie. Centrally, zygomatic synchrony in both emotional contexts predicted subsequently reported affiliative feelings of dyad members. These results demonstrate that a naturally unfolding smiling behavior is highly contagious. They further suggest that zygomatic synchrony functions as a social facilitator, eliciting affiliation towards previously unknown others.

## Introduction

Connecting with others is a fundamental human drive ^1^, arising both in the context of close relationships and towards complete strangers. Given its central role in human flourishing and well-being ^2^, it is important to understand the biologically-wired mechanisms which allow people to rapidly and effortlessly form social bonds. It has been suggested that one such mechanism is interpersonal synchrony - the pervasive tendency of individuals to become coupled or synchronized with each other during interaction^3–8^. Past research has indeed suggested that interpersonal alignment of neural responses between interacting individuals binds them into a social unit, promoting mutual emotional and social states^3,6–9^. However, synchrony in the humans’ main communication channel, i.e. facial dynamics, has not been systematically studied.

Using hyper-scanning methodology and measures collected over time, research has demonstrated that verbal and non-verbal social interactions involve temporally coordinated autonomic and neural responses between people^4,10,11^. Such synchrony is grounded in multiple exchanges of sensory signals between individuals, which allow for continuous reciprocal adaptations to the incoming social input ^6,10,12^. Accordingly, neural and autonomic synchronies, which unfold in the internal milieu, are intricately connected with externally exchanged information. Facial displays serve as one of the most pronounced external responses unfolding directly in a social environment. Moreover, facial signals play a profound social role, as they do not only express emotions but also convey intentions to others^13–16^. As such, the social spread of facial dynamics may bridge the gap between individual behaviors, physiological synchrony and higher-order emotional-social phenomena, such as shared experiences and affiliative feelings.

In the current research, we aim to assess spontaneous synchrony in facial dynamics of co-present individuals and to study its emotional and social function. We focus on synchrony in positive facial signals, that is a smiling behavior, for several reasons. In contrast to computerized paradigms, in a naturalistic social context subjects’ attention is not pre-fixed on targets’ faces. Positive facial expressions seem to be beneficial in such a condition, as they are characterized by a distinctive visual saliency and diagnostic value, even when presented in a peripheral visual space ^17–19^. In addition, the temporal dynamics of positive facial signals can be reliably assessed with continuous electromyography measures (EMG) over zygomaticus major facial muscle^20–25^. Finally, while social mimicry of negative facial expressions is dependent on multiple social factors^26^, mimicry of positive facial responses is a more robust phenomenon ^20,27,28^. Thus, interpersonal synchrony in zygomatic responses promises to be a valuable measure to study how visually salient facial behaviors are reciprocated between people and across time.

Humans’ propensity to react with corresponding facial expressions to facial displays of others has been massively documented by the facial mimicry research ^26,28–34^. Specifically, studies have demonstrated that happy faces, presented via pictures or short video sequences, elicit zygomatic activation in observers ^29,30,33,35–37^. While this research provides a solid basis for hypothesizing an existence of social coordination in zygomatic dynamics, it has been criticized on two main grounds. First, zygomatic activation in observers is triggered not only by targets’ happy faces, but also by joyful voices or body postures^38,39^. Accordingly, zygomatic activation in computerized paradigms cannot be taken as a direct evidence of interpersonal coordination of facial muscles, as it may stem from positive emotions elicited by targets’ smiles^34,40^. Second, computerized paradigms are devoid of social context, hence suffering from poor ecological validity and impoverished ability to assess the social outcomes of such coordination ^20,26,41^. Beyond mimicry, research has also tapped into smiling reciprocity – that is a social behavior of returning a smile of your interaction partner. Employing facial coding^21,42^ and EMG measures^20^, a few studies have demonstrated that people return their partners’ smiles with high probability during both positive and negative conversations. While these studies have provided a more direct evidence for interpersonal coordination of facial signals, they relied on explicit verbal interactions. A face-to-face communication encompasses complex appraisal processes, such as social expectations and communicative motivations, hence hampering the ability to assess relatively automatic spontaneous synchrony in facial dynamics as well as the outcomes of such synchrony.

To investigate spontaneous facial synchrony, while preserving naturalistic social dynamics, we employed a well-established co-viewing paradigm, in which dyads of participants viewed emotional movies together without directly interacting ^3,43–46^. In this paradigm emotional movies serve as catalyzers of emotional behaviors, eliciting non-verbal emotional signals from participants. The lack of direct communication between participants as well as a real-life like social context of co-viewing reduces the impact of high-order cognitive processes. Notably, although participants’ attention is directed towards an external stimulus, they still act as social agents, free to modify their covert and overt attention and to form social impressions^44^. Accordingly, this paradigm has been previously shown to elicit profound interpersonal processes in co-viewing dyad members, as evidenced in their emotional^44^ and physiological^47^ responses.

We assessed zygomatic EMG and cardiovascular signals in previously unacquainted dyads of participants who co-viewed a positive and a negative movie and during a non-emotional baseline. Participants provided emotional ratings following each movie and reported on their affiliative feelings towards each other at the end of the experiment. Zygomatic and cardiovascular synchrony were quantified both in co-viewing dyad members (hereafter, real dyads) and in randomly shuffled control dyads (members of different dyads who viewed the same movies but not together; hereafter, random dyads). Synchrony in random dyads represents the *movie-driven effects,* which manifest similarity in the affective dynamics, elicited by the movies^47,48^. Synchrony in real dyads incorporates *dyad-specific interpersonal processes* unfolding at a dyad-unique timeline. Given the well documented specificity of zygomatic activity to positive emotional experiences, we expected to find enhanced zygomatic synchrony during the positive as compared to the negative movie or baseline, both in real and in random dyads. Centrally to the present study, we hypothesized that zygomatic synchrony will be enhanced in real dyads as compared to random dyads, signifying dyad-specific interpersonal coordination of facial signals. Previous studies have reported that in naturalistic social contexts individuals frequently exhibit zygomatic activity during both positive and negative emotional experiences ^20,46^. Accordingly, we compared the real to random dyads synchrony across all experimental conditions.

To elucidate the role played by zygomatic synchrony in emotional transmissions within real dyads, we studied its association with cardiovascular synchrony and convergence in feelings. Ample studies have demonstrated that facial mimicry may lead to emotional contagion ^49^. In addition, previous research has suggested that contagious emotional transmissions involve synchronization of autonomic signals^3,50^. We hypothesized that zygomatic synchrony is involved in contagious transmissions of positive emotional signals. Accordingly, we expected to find tight links between zygomatic synchrony, cardiovascular synchrony and emotional convergence during a positive but not during a negative movie.

Finally, we examined the affiliative social role of zygomatic synchrony. Smiling is perceived as an affiliative signal^51,52^. Furthermore, social motives towards the other person were found to impact social smiling^44,45^. However, whether spontaneous zygomatic synchrony directly facilitates social connections is currently unknown. To test this hypothesis, we examined whether zygomatic synchrony in previously unacquainted participants is associated with their subsequently reported affiliative feelings towards each other.

## Methods

### Participants

26 dyads of female students (age, M=23±2.94) participated in the study for academic credits, after signing a written informed consent. Female participants were used to enhance the probability for social smiling^53,54^ and to avoid gender-specific differences in response to movies. As validated prior to arrival, dyad members were not acquainted with each other. Experimental procedures were approved by the Ethics Committee for Behavioral Studies at the Interdisciplinary Center Herzliya. The methods were carried out in accordance with the approved guidelines.

### Materials and Procedure

Dyads of previously unacquainted participants arrived to the lab and were told that the aim of the experiment is to measure physiological responses to movies. They were seated side by side on two adjacent armchairs in front of a 20-inch computer screen. The viewing distance to presentation screen was approximately 70 cm. Participants were connected to physiological sensors (see below) and were asked to refrain from talking and making gross movements throughout the entire experiment. Baseline measures were recorded for five minutes while participants sat silently and calmly and kept their eyes open. After the baseline, participants took part in a co-viewing experiment which is not reported here and then viewed a positive and a negative movie. Two movie excerpts, used in previous studies ^3,23,47^, were taken from the horror film “*Paranormal Activity”* (394s) and the popular entertainment TV show “*Britain’s Got Talent”* (364s) to elicit negative and positive emotions, respectively. Each emotional movie was preceded by a ten second countdown screen and the order of the movies was counterbalanced across participants. After each movie, participants rated their emotional responses to the movies. At the end of the experiment, participants filled out an affiliation questionnaire in different rooms.

### Measures

#### Subjective reports

##### Emotional experience

Following each movie, participants rated the intensity of the positive and the negative feelings elicited by the movie, using two unipolar 9-point Likert scales.

##### Affiliative feelings

Participants rated the degree to which they felt closeness, similarity and rapport towards their co-present partner, on a 9-points Likert scale. Items showed Cronbach’s alpha = 0.89, and were averaged to form an *affiliation* score.

##### Video coding of smiling

To validate that zygomatic signals are associated with smiling behavior, a research assistant with an extensive previous expertise in facial coding, coded frequency of smiling events from video recordings of participants. Frequency of individual smiles and frequency of shared (co-occurring) smiles were coded separately.

##### Physiological data collection and preprocessing

Continuous physiological measures were recorded with a Bionomadix (BIOPAC Systems Inc., Santa Barbara, CA) MP150 data acquisition system and Biopac^®^ AcqKnowledge 4.4 software at 1000Hz sampling rate.

Cardiovascular activity was recorded using a modified lead II configuration. The heart period series (inter-beat interval; IBI) were assessed using the Mindware HRV 2.16 biosignal processing module (Mindware, Ohio). This procedure included: (a) identifying the R–R intervals; and (b) detecting physiologically improbable R–R intervals based on the overall R–R distribution using a validated algorithm ^55^. Afterwards, data were also inspected manually to ensure that R-waves were correctly identified. Data that included more than 10% undefinable Rs was excluded from the analysis. Lastly, IBI series were transformed to continuous 1 Hz heart rate time-series (HR) using custom software.

Facial electromyography (EMG) was collected over the zygomaticus major and corrugator supercilii sites, placed at the left side of the face ^56^. This paper focuses on zygomatic responses and thus reports the results for the zygomatic (EMGZYG) signals only. EMG signals were filtered with a notch 50 Hz filter. Manual inspection for artifacts was made using custom software ^57^. Frequency analysis of EMG data has been applied, in which a Fast Fourier Transform was performed on all artifact-free data to derive estimates of spectral power density (*μV*) in the 45–200-Hz frequency band in one sec windows^24,25,58^. Finally, these values were log-transformed to normalize the data. This analysis yielded continuous 1 HZ EMGZYG time-series. The first 20 seconds and last 5 seconds were cropped from the resulting HR and EMGZYG time-series to remove non-specific accommodation effects and preprocessing artifacts. For the mean level analysis, we quantified mean EMG responses by averaging the sample points across time during each experimental condition. To assess the effects of the movies on mean EMG activity, the EMG scores of the dyad members were averaged^59^, and within-subjects ANOVA was computed on these dyadic scores.

##### Missing data

Dyads in which one or both dyad members failed to refrain from talking were excluded from analysis. This resulted in an exclusion of five, one and three dyads from the baseline period, the positive movie and the negative movie conditions, respectively. At the preprocessing stage, data was monitored for low signal quality and for excessive artifacts, such as multiple non-specific spikes or motion artifacts. Exclusion of one participant has led to an exclusion of the dyad. Overall, the percent of missing data points for each analyzed channel comprised 12% for HR and 10% for EMGZYG.

### Data Analysis

#### Assessment of synchrony

To assess synchrony we computed a cross-correlation between two individual z-normalized physiological response time-courses within ±3 seconds temporal lag^3,47^. Maximal correlation values, as well as temporal lag at which these values occurred were extracted. For the parametric statistical analysis of synchrony, the maximal correlation values were Fisher z-transformed.

#### Comparing zygomatic synchrony in real dyads and in random dyads

The central result of this study was derived from comparing the zygomatic synchrony computed for real dyads (participants who viewed the movies together) to zygomatic synchrony computed for random dyads (participants from different dyads who viewed the same movies but not together)^3,60^. We hypothesized that participants who viewed the movies together would exhibit higher levels of zygomatic synchrony relative to those who viewed the same movie but in randomly shuffled dyads. To test our hypothesis, we computed zygomatic synchrony in all possible random dyads in the sample and constructed empirical control sampling distribution of random dyads’ synchrony, using bootstrapping procedure with 1000 iterations. The statistical likelihood of zygomatic synchrony in the real dyads was assessed directly from this control distribution (Supplementary Figure 1). This procedure was done separately for each experimental condition (baseline, negative, positive). A similar approach was taken with the cardiovascular measures, yielding estimates of HR synchrony in real and in random dyads.

#### Association with self-reports

We conducted correlation analyses between indexes of cardiovascular and zygomatic synchrony and self-reports, using the Spearman rank correlation. Dyad level self-reports were computed using following guidelines: 1) emotional convergence was computed as the sum of dyad members’ emotional ratings divided by the absolute difference between the dyad members’ emotional ratings (*E*_1_ + *E*_2_)/(*abs*(*E*_1_ − *E*_2_) + 1). Emotional convergence was assessed for positive feelings during the positive movie and for negative feelings during the negative movie 2) Dyads’ affiliation scores were computed by averaging dyad members’ ratings, as recommended for dyadic designs ^59^.

### Data availability

The datasets generated and analyzed during the current study are available from the corresponding author on reasonable request.

## Results

### Emotional movies elicited differential affective and zygomatic response patterns

To begin with, we validated that the positive and the negative emotional movies elicited the expected emotional and facial response patterns. Repeated measures ANOVA with type of movie (positive, negative) and valence (positive, negative) factors on self-reports showed the predicted valence by movie interaction (F(1,18)=726, p<0.001). A high level of positive feelings (M=7.23±1.03) and a negligible level of negative feelings (M=0.6±1.07) was reported for the positive movie. A high level of negative feelings (M=6.75±0.76) and a negligible level of positive feelings (M=0.77±0.89) was reported for the negative movie. An expected pattern was obtained also for the mean-level zygomatic EMG, which exhibited higher activity during the positive compared to the negative movie (t(18)=11.871, p<0.001). Zygomatic activity during the negative movie was higher than the one during baseline, at marginally significant level (t(13)=2.009, p<0.066).

### Zygomatic synchrony in real and random dyads

Central to the present study, we asked whether co-present participants became spontaneously synchronized with each other in their zygomatic facial activity and whether this synchrony was dyad-specific. Figure 1A presents an example of zygomatic activity in two different dyads during the positive movie. As can be seen in the figure, the co-viewing dyad members showed clear temporal alignment of zygomatic dynamics with each other. Moreover, each dyad showed idiosyncratic zygomatic response patterns, leading to higher within-dyad [0.63, 0.66] than across-dyads correlations [0.45, 0.46, 0.39, 0.45].

We computed zygomatic synchrony in real dyads and compared it to zygomatic synchrony in random dyads. Figure 1B presents the mean zygomatic synchrony in real and random dyads in all experimental conditions. The descriptive statistics for the zygomatic synchrony in real and random dyads are presented in Table 1. As evident in Figure 1B, zygomatic synchrony was enhanced during the positive movie compared to the negative movie and baseline. This difference occurred both for the real dyads (repeated measures ANOVA (F(2,26)=8.15, p<0.001; positive vs negative, t(18)=2.9, p<0.008) and for the random dyads (Wilcoxon signed rank test: positive vs negative, z=23.56, p<0.001). Significantly enhanced zygomatic synchrony during the positive movie comes in line with the previously reported zygomatic specificity for positive emotions ^22,61^. It reflects the consistent modulations of zygomatic activity exerted by the positive emotional timeline, which was shared across all participants.

**Figure 1.**
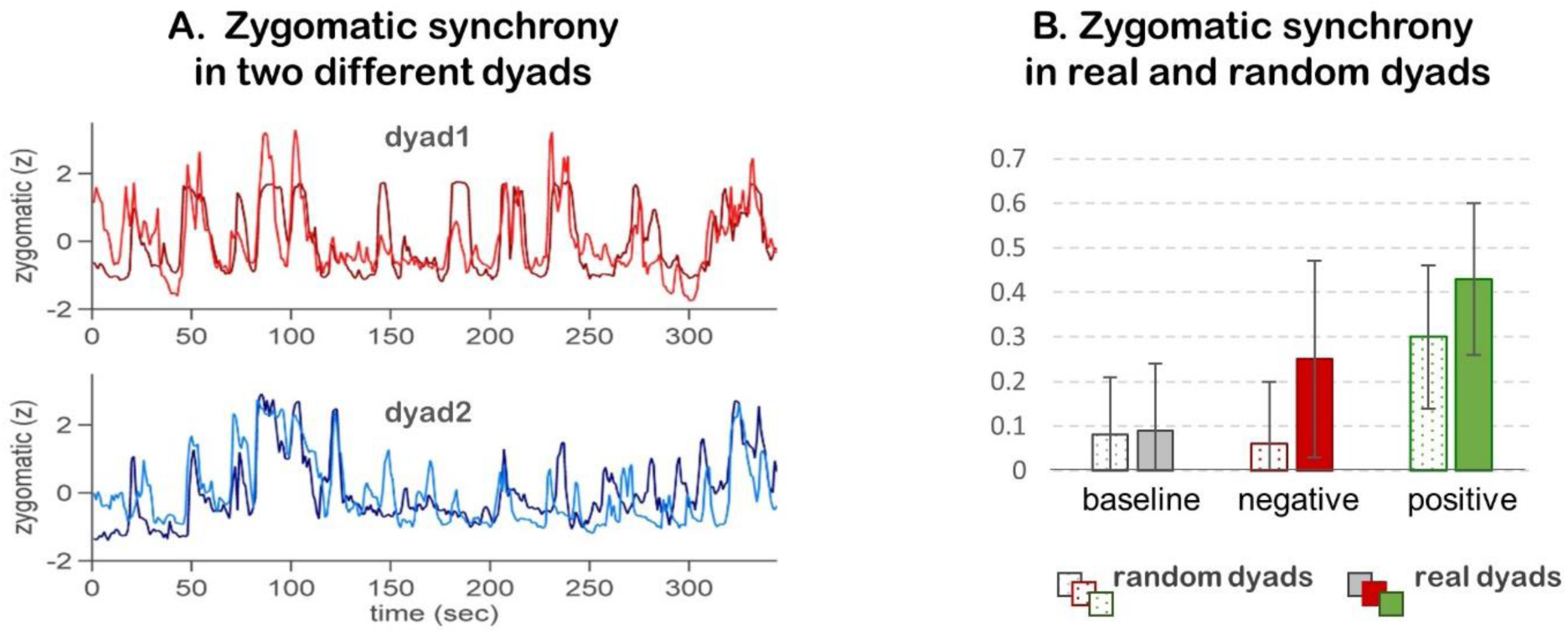
A. Zygomatic synchrony in two different dyads. Zygomatic EMG responses in two different co-viewing dyads (dyad1-red colors, dyad 2 – blue colors) are presented. **B. Zygomatic synchrony in real and random dyads.** Zygomatic synchrony was assessed in real dyads (i.e. participants who viewed the movies together, marked by full bars) and in random dyads (i.e. participants who viewed the same movies but not together, marked by patterned bars), during non-emotional baseline (grey), the negative movie (red) and the positive movie (green) conditions.

We compared the zygomatic synchrony in real and random dyads, using a non-parametric bootstrapping approach (Methods, Supplementary Figure 1A). In line with our central hypothesis, zygomatic synchrony in real dyads was significantly higher than the one in random dyads, demonstrating dyad-specific contagious zygomatic transmissions. The enhanced zygomatic synchrony in real dyads was evident during both the positive and the negative movies but not during non-emotional baseline.

To validate that zygomatic synchrony was indeed driven by smiling behavior, we coded individual and shared smiles from the video-recordings of participants during the movie viewing (Supplementary Figure 2). We found that although the positive movie elicited significantly higher amount of smiles (t(19)=5.3, p<0.001), participants smiled both during the positive (M=19.1±9) and the negative (M=7.8±7.3) movies, and some proportion of these smiles was shared (positive: M=5.3±4.2; negative: M=0.9±4.9). Notably, frequency of shared smiles was correlated with zygomatic synchrony during the positive (ρ=0.54, p<0.014) and the negative (ρ=0.526, p<0.021) movies.

To further understand the nature of zygomatic synchrony during the positive and the negative movies, we compared the temporal lags at which co-viewing participants’ zygomatic responses exhibited maximal correlation in these two emotional contexts. This analysis showed that the dyad members’ zygomatic responses were more tightly linked (i.e. shorter temporal lags) during the positive (M=0.48±0.14 sec) as compared to the negative (M=1.75±0.24 sec) movies (t(1,18)=4.65,p<0.001).

**Table 1.** Descriptive statistics for the zygomatic and cardiovascular synchrony indexes in real and in random dyads. Table presents means, standard deviations and range of synchrony indexes in real dyads; as well as means and standard deviations of control sampling distributions of synchrony in random dyads. It also presents the p values for the non-parametric test of hypothesis that synchrony in real dyads is larger than synchrony random dyads.

|  | ZYGOMATIC SYNCHRONY |  |  | CARDIOVASCULAR SYNCHRONY |  |  |
| --- | --- | --- | --- | --- | --- | --- |
| | real dyads | random dyads | $p$ | real dyads | random dyads | $p$ |
| baseline | $0.09\pm 0.16$ [0.58] | $0.08\pm 0.021$ | $p=0.24$ | $0.07\pm 0.12$ [0.49] | $0.06\pm 0.017$ | $p=0.24$ |
| negative | $0.24\pm 0.22$ [0.92] | $0.06\pm 0.021$ | $p<0.001$ | $0.15\pm 0.11$ [0.43] | $0.13\pm 0.017$ | $p=0.054$ |
| positive | $0.44\pm 0.17$[0.58] | $0.3\pm 0.023$ | $p<0.001$ | $0.17\pm 0.17$ [0.68] | $0.12\pm 0.014$ | $p<0.001$ |

Taken together, these results demonstrate that co-viewing participants exhibited robust synchronization in their zygomatic responses while viewing emotional movies but not during non-emotional baseline (Table 1, Supplementary Figure 1). Such synchrony represented direct, dyad-specific facial coordination, as it was higher in real as compared to random dyads and correlated with the amount of shared smiling. While statistically robust zygomatic synchrony was observed in both emotional contexts, participants’ zygomatic responses were more similar in their dynamics and more tightly linked in time during the positive as compared to the negative movie.

### Association with cardiovascular synchrony

To investigate whether zygomatic synchrony is linked with synchronization of visceral autonomic responses, we examined its association with synchrony in participants’ heart rates. First, we assessed cardiovascular synchrony in real and random dyads (Table 1). Both real and random dyads exhibited higher cardiovascular synchrony during emotional movies as compared to baseline (random dyads: negative vs baseline: z=8.39, p<0.001; positive vs baseline: z=9.12, p<0.001; real dyads: positive vs baseline, t(16)=2.46, p<0.05; negative vs baseline, t(14)=1.9, p=0.06). In contrast to zygomatic synchrony, cardiovascular synchrony wasn’t significantly different for the positive and negative movies (Wilcoxon signed rank test, z=1.1, p>0.25; t(18)=0.5, p>0.25). This pattern of results supports the established links of cardiovascular responses with emotional processing ^3,61,62^. Centrally to the present paper, real dyad members exhibited higher synchrony in their heart rates as compared to random dyads. Such dyad-specific synchrony was significant during the positive (p<0.001) and marginally significant during the negative (p=0.054) movie but not during baseline (p>0.25) (Table1, Supplementary Figure 1B).

We next asked whether the degree of a dyad’s zygomatic synchrony was associated with this dyad’s cardiovascular synchrony. For that aim, we computed Pearson correlations between zygomatic and cardiovascular synchrony indexes across dyads. We found that synchrony indexes were significantly correlated during the positive movie (r=0.448, p<0.05). Such association didn’t occur during the negative movie (r=0.05). In other words, during the positive but not during the negative emotional experience, synchronization of zygomatic responses in dyad members was intricately related to the unfolding of mutual autonomic states.

### Association with emotional convergence

The above results demonstrated that co-viewing participants exhibited idiosyncratically synchronized zygomatic and cardiovascular responses during both emotional movies but not during non-emotional baseline. To elucidate the role of zygomatic and cardiovascular synchronies in contagious emotional transmissions across dyad members, we calculated correlations between indexes of zygomatic and cardiovascular synchrony and indexes of emotional convergence. The results showed that cardiovascular synchrony was positively associated with emotional convergence in both positive (ρ =0.51, p<0.01) and negative (ρ =0.67, p<0.001) feelings. In contrast, zygomatic synchrony was positively related with emotional convergence in positive (ρ =0.462, p<0.02) but not in negative (ρ =0.02, p>0.25) feelings. Taken together, these findings indicate that during the positive movie both facial and cardiovascular synchronies were involved in contagious transmissions of positive emotional signals between participants. In contrast, during the negative movie zygomatic and cardiovascular synchrony seemed to play different functions, as only the cardiovascular but not the zygomatic synchrony was linked with a shared negative experience.

It could be suggested that the links between synchrony and emotional convergence in dyad members are not driven by direct interpersonal transmissions, but rather by the similarity of their emotional responses. Accordingly, participants who similarly reported high positive feelings could also show more similar facial or heart rate response patterns during the viewing. To control for such possibility, we assessed the association between zygomatic synchrony and emotional convergence in random dyads, in which interpersonal transmissions couldn’t occur. No substantial positive correlation was observed (ρ =-0.01) for zygomatic synchrony and emotional convergence in random dyads during the positive movie. Similar results were obtained for the cardiovascular synchrony during the positive (ρ =0.03) or negative (ρ =0.04) movies.

### Association with affiliative feelings

Finally, we examined the affiliative social role of zygomatic synchrony. At the end of the experiment participants reported the degree of their affiliative feelings towards the other participant. We computed correlations between affiliation scores and synchrony indexes during the first and the last presented movies (Figure 2). To account for the inherent differences in the mean level of synchrony during the positive and the negative movies, we mean-normalized the synchrony indexes within each emotional condition. Results showed that zygomatic synchrony in dyad members during the last (ρ=0.739, *p*<0.001) but not during the first movie (ρ=-0.04, *p>*0.25) were highly correlated with their post experiment affiliative feelings towards each other. We further validated this analysis by assessing correlations for the positive and the negative movies in separate. Both the positive and the negative movies, when presented last, were associated with affiliation scores (all *p*s < 0.009). In contrast, first-presented movies didn’t show such associations (all *p*s > 0.35). Taken together, these results suggest that zygomatic synchrony elicited affiliative feelings between previously unacquainted participants. The lack of association with affiliative feelings during the first movie speaks against the alternative possibility that zygomatic synchrony reflected initial social perceptions.

**Figure 2.**
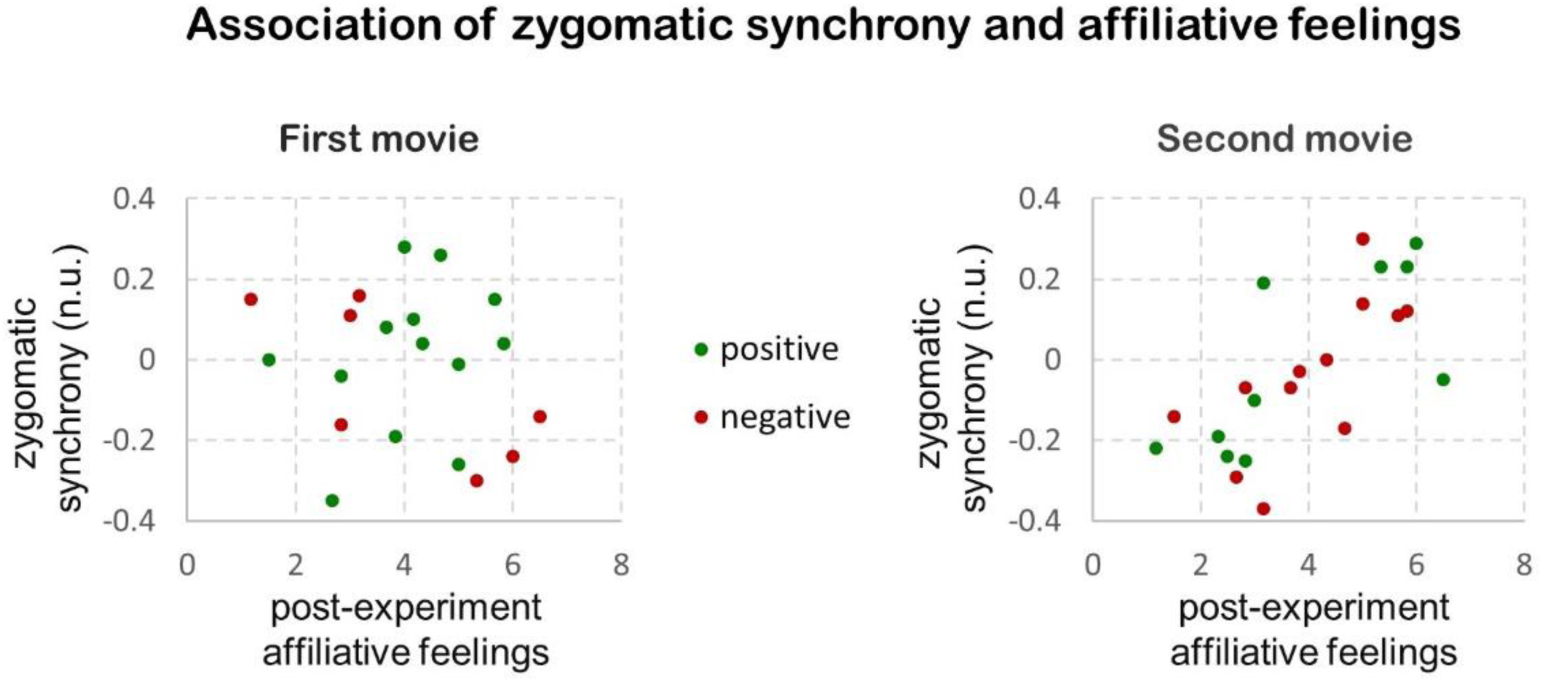
Association of zygomatic synchrony and affiliative feelings. Zygomatic synchrony during the last (ρ=0.739), but not during the first (ρ=-0.04) presented movie was correlated with post-experiment affiliative feelings of co-viewing participants. Axis y represents zygomatic synchrony indexes, mean-normalized within each emotional condition. Scores arriving from the positive and the negative movies are marked with green and red colors, respectively. *n.u. normalized units.*

## Discussion

In this research, we examined a spontaneous social synchrony in zygomatic responses of co-viewing participants, as well as its role in emotional and affiliative processes. We measured moment-by-moment changes of zygomatic and cardiovascular responses in dyads of previously unacquainted female participants while they co-viewed a positive and a negative movie and during a non-emotional baseline. To index dyad-specific processes as opposed to stimulus-driven processes we computed synchrony in real dyads who viewed the movies together and compared it to synchrony in random dyads, who viewed the same movies but not together.

The main result of our study demonstrates that real dyad members exhibited robust synchrony in their zygomatic responses. Although all participants shared similar affective dynamics, elicited by the movies, co-viewing dyad members were synchronized in their zygomatic responses above and beyond this movie-driven similarity, as evidenced by real to random dyads comparison. This result suggests that smiling dynamics of co-present others are highly contagious. It significantly extends previous research on automatic mimicry of smiles^28–30,37,63,64^ to continuous interpersonal coordination of facial dynamics unfolding in a naturalistic social context. The pervasiveness of observed zygomatic synchrony in non-acquainted individuals who did not have explicit social goals, supports an involvement of automatic, bottom-up processes^29,37^ driving individuals to couple with others in their facial responses.

Curiously, we found that zygomatic synchrony in real dyads emerged during both the positive and the negative emotional contexts (but not during non-emotional baseline). More specifically, while the overall intensity of zygomatic synchrony during the negative movie was diminished as compared to the positive movie, it was nevertheless statistically robust in the co-viewing dyad members. Video analysis of participants’ faces, confirmed that zygomatic synchrony during both movies was linked with shared smiles. This finding is further corroborated by previous studies which demonstrated that people tend to smile in negative social contexts, such as when co-viewing a sad movie with a friend^46^ or during negative social interactions^20^. Building upon previous models of interpersonal emotional processes^14,65,66^, it can be speculated that co-viewing a negative movie with a stranger might violate socially accepted norms and elicit complex feelings and appraisals. Accordingly, zygomatic synchrony in the negative movie can represent the result of such appraisals, signifying social intents and shared understandings of the social-emotional situation ^44,67^. In support of such interpretation, we found that the temporal lags between participants’ zygomatic responses were significantly longer during the negative as compared to the positive movie, potentially reflecting high-order appraisal processes. While this wasn’t the main focus of the current study, this result suggests that zygomatic synchrony can occur both via automatic bottom-up processes when appearing in a congruent emotional context and via higher order social appraisals in an emotionally in-congruent context. Future studies are needed to systematically examine the effects of social appraisals on facial synchrony.

Human face is a profound communicative device equally connected with internal responses and experiences and external behaviors unfolding in a social space. Here we examined interpersonal synchrony as a multi-system phenomenon, by assessing the association of zygomatic synchrony with convergence in cardiovascular responses and subjective experiences. First, we found that synchrony in co-viewing dyad members wasn’t reserved for facial dynamics, but occurred in their autonomic responses as well. Specifically, dyad members were significantly synchronized in their cardiovascular responses during both emotional movies, but not during baseline. Such spontaneous synchrony in autonomic responses was previously demonstrated for friends^3^ and here extended for previously unacquainted dyad members. We also found that cardiovascular synchrony was associated with emotional convergence in both positive and negative feelings, providing strong support for its central role in contagious emotional transmissions between individuals^3^. As for its association with zygomatic synchrony, it differed substantially in different emotional contexts. During the positive movie zygomatic synchrony significantly co-varied with cardiovascular synchrony as both manifested convergence in positive feelings. This finding is in line with previous research linking synchronous physiological processes in interacting individuals with shared emotional experiences, such as emotional contagion and empathy ^3,7,68,69^. In contrast, during the negative movie, zygomatic synchrony didn’t show such association with cardiovascular synchrony or with emotional convergence, suggesting that it played a different functional role.

Finally, our results demonstrated that spontaneous zygomatic synchrony had a profound social impact on co-viewing strangers, enhancing their affiliative feelings towards each other. Multiple previous studies have shown that behavioral mimicry leads to enhanced liking ^70–72^. As for the case of facial mimicry, previous research has demonstrated that smiling mimicry is influenced by the social motives of observers ^44,45,73^ and by the social status of the mimicry targets^27,28,45,64^. The naturalistic social nature of the current study enabled us to demonstrate that zygomatic synchrony serves as an affiliation building behavior, enhancing social bonds in previously unacquainted participants. In particular, we found that zygomatic synchrony predicted the subsequently reported affiliative feelings of participants and that such links existed for zygomatic synchrony in both positive and negative emotional contexts. While it could be suggested that links with affiliation were mediated by initial social impressions of participants or their social motivations, our results speak against this possibility. Specifically, we found that the association of zygomatic synchrony with post-experiment affiliation ratings was observed only for the last but not for the first part of the experiment. Extending previous models, which assign a unique affiliative value to smiling^13,15,52^, our findings suggest that synchrony in smiling dynamics is a profound social behavior which builds affiliative context between people.

## Conclusion

Using a naturalistic social paradigm and physiological measures assessed over time this study demonstrated that previously unacquainted individuals exhibit robust synchronization in their smiling dynamics when co-viewing movies together. Centrally, such zygomatic synchrony was evidenced above and beyond the movie-driven effects and reflected contagious facial transmissions in co-viewing dyad members. In a positive emotional context zygomatic synchrony was linked with cardiovascular synchrony as well as with convergence in positive feelings. The robust occurrence of zygomatic synchrony in previously unacquainted participants lacking explicit social goals or direct communication speaks to the involvement of automatic, bottom-up processes leading people to dynamically coordinate their smiling behaviors with co-present others. Zygomatic synchrony was observed in a negative emotional context as well. However, it wasn’t associated with cardiovascular synchrony or with positive feelings, and occurred at longer temporal lags, suggesting an involvement of higher-order appraisal processes. Centrally, zygomatic synchrony in both emotional contexts predicted subsequently reported affiliative feelings of participants towards each other. Taken together, our findings demonstrate that smiling dynamics have a high probability to spread in a social environment, eliciting mutual experiences and affiliative feelings.

## Acknowledgements

This work was funded by Israel Science Foundation (ISF) Grant 698/15 to YG. We thank Roni Nadav for a tremendous help in data collection and preprocessing. We thank Adam Hakim and Tali Aloni for providing help with data analysis. We thank Alexandra Kopelansky for coding facial expressions.

## Authors contributions

YG, NLB, DM. designed the experiment. DM collected the data and helped with the analysis. YG and NLB designed the analysis. YG wrote the manuscript, NLB and DM provided critical review. All authors approved the manuscript.

## Competing Interests

The authors declare no competing interests.

